# Cellular Mechanisms Underlying Melanoma Brain Metastasis

**DOI:** 10.1101/2025.11.24.689301

**Authors:** Bengi Ruken Yavuz, Hyunbum Jang, Ruth Nussinov

**Affiliations:** Cancer Innovation Laboratory, National Cancer Institute, Frederick, MD 21702, USA; Computational Structural Biology Section, Frederick National Laboratory for Cancer Research, Frederick, MD 21702, USA; Department of Human Molecular Genetics and Biochemistry, Sackler School of Medicine, Tel Aviv University, Tel Aviv 69978, Israel

**Keywords:** melanoma brain metastasis, molecular mechanism, combination drugs, neurotrophin signaling, PI3K/AKT pathway, MAPK pathway

## Abstract

Brain metastasis is a major contributor to melanoma-related mortality, arising from both genomic alterations and dynamic adaptations to the brain microenvironment. Tumor cells must cross the blood–brain barrier and exploit brain-specific signaling and metabolic landscapes to establish metastases. Brain metastases display molecular and phenotypic programs distinct from other organs, reflecting pre-existing and acquired mutations during progression. We analyzed *primary melanoma, metastatic melanoma, and metastatic brain tumors* genomic datasets to identify mutated proteins and signaling pathways that are common to or distinct among these tumor types. Our comprehensive computational analyses uncovered key pathways and proteins associated with brain metastasis. Brain metastases exhibit unique mutational landscapes and convergent alterations in neurotrophin, MAPK, and PI3K/AKT signaling compared with primary and metastatic melanoma. Proteins especially frequent in brain metastases pathways support survival, proliferation, and adaptation in the brain microenvironment. Pharmacology-wise, rational strategies such as combining c-KIT and *VEGFR* inhibitors can suppress converging oncogenic pathways and mitigate resistance in advanced melanoma. Co-targeting neurotrophin receptors with MAPK pathway members, particularly in combination with *BRAF* inhibitors, appears a promising approach for treating melanoma brain metastases. *Our findings underscore the molecular heterogeneity of melanoma brain metastases, shaped by interconnected signaling networks*, *and reveal therapeutic targets that may diverge from those in other metastatic sites*.

## Introduction

Metastasis, the spread of cancer cells from a primary tumor to distant organs, is the leading cause of cancer-related mortality [1–3]. This dissemination is not random but demonstrates *organotropism*, wherein specific cancers preferentially colonize distinct organs [4–6]. Prostate cancer tends to metastasize to the bone; melanoma to the liver and brain; and breast cancer to the bone, lung, liver, and brain, with luminal subtypes particularly favoring the bone [7]. These patterns have been consistently confirmed in large-scale clinical studies across cancer types [6–8]. A recent review of melanoma summarized its aggressive nature, key genetic drivers, and evolving therapeutic landscape [9]. While targeted therapies and immune checkpoint inhibitors have improved outcomes, major challenges persist, including therapy resistance and recurrence [10].

In melanoma, brain metastasis is associated with distinct molecular features not always present in extracranial, or other body sites. For example, some *KRAS* mutations which are rare in melanoma but enriched in early brain metastases and exclusive with other MAPK mutations were identified (e.g., *BRAF^V600^*, *NRAS*) [11]. These *KRAS*-mutant clones appeared to originate early, suggesting a brain-specific dissemination route. Gene expression analysis of intracranial and patient-matched extracranial melanoma metastases revealed site-specific molecular programs [12]. Intracranial lesions showed downregulation of immune-related genes and activation of brain-like transcriptional signatures, with frequent alterations in cytokine–receptor interaction, calcium signaling, ECM–receptor interaction, cAMP signaling, Jak/STAT, and PI3K/AKT pathways.

Hyperactivation of the PI3K/AKT pathway is a recurrent finding in melanoma brain metastases. It is also critical for the brain, where it regulates complex neuronal functions such as synaptic plasticity, memory, and neurogenesis [13, 14], making it especially receptive and conducive to melanoma metastases and possibly one reason for its thriving there. In preclinical models, the PI3K inhibitor buparlisib reduced *AKT* activity, inhibited tumor growth, and induced apoptosis in melanoma cells in both in vitro and intracranial mouse models, irrespective of *BRAF* or *NRAS* status [15]. These results support PI3K inhibition as a promising therapeutic strategy, currently under clinical investigation. Redmer et al. [16] highlighted the multifactorial nature of melanoma brain metastases, implicating not only tumor-intrinsic mutations but also dynamic interactions with the brain microenvironment. In line with this, key mediators include PI3K/AKT, as well as STAT3, and TGFβ signaling, both also critically important for the brain, where they regulate a wide range of essential functions [17], along with molecules such as *PLEKHA5*, *CD271*, and *CCR4*, which facilitate blood–brain barrier transmigration, survival, and colonization [18–20]. Notably, the brain is also exceptionally conducive to breast cancer, which is also often marked by overactive PI3K/AKT, as is lung cancer, both frequently metastasizing into the brain. Both tumors also manifest strong STAT3, and TGFβ signaling [21–24], sharing signaling properties of melanoma, and the hosting brain.

High-resolution genomic analyses underscore the brain-specific nature of certain melanoma metastases. Targeted sequencing of 29 driver genes in matched intracranial and extracranial metastases revealed brain-exclusive mutations in 11 of 16 patients [25]. Mutations in chromatin remodelers such as *ARID1A*, *ARID2*, *SMARCA4*, and *BAP1* were found only in brain metastases, supporting a branched evolution model and offering potential targets for brain-directed therapies. Multi-omics profiling has delineated two predominant phenotypic states in melanoma metastases [26]: one associated with *E-cadherin* expression, indicative of a proliferative and pigmented state, and the other characterized by *NGFR* expression, associated with invasiveness, stemness, and therapy resistance. A phenotypic shift from E-cadherin to *NGFR* marks disease progression and drug resistance, with notable differences between *BRAF*-mutant and wild-type melanoma brain metastasis. Epigenetic alterations, alongside genomic changes, play a pivotal role in melanoma brain metastasis. Promoter methylation and gene expression profiling revealed three patient subgroups in matched brain and extracranial metastases [27].

Single-cell and spatial transcriptomic analyses have provided further granularity. Treatment-naive melanoma brain metastasis shows increased chromosomal instability, neural-like transcriptional programs, and distinct metabolic and immune evasion strategies compared to extracranial lesions [28]. A comprehensive single-cell atlas of 108 human brain metastases across multiple cancer types highlighted conserved hallmarks of brain metastasis: chromosomal instability, neural-like transcriptional signatures, and a microenvironment dominated by immunosuppressive myeloid and stromal cells [29]. Using a multi-omics approach integrating spatial transcriptomics with multi-region exome, proteome, and transcriptome profiling in four patient tumors, the study revealed patient-specific immune infiltration patterns and protein-level heterogeneity [30].

Brain metastasis requires that circulating tumor cells (CTCs) first traverse the blood–brain barrier (BBB)—a specialized structure composed of endothelial cells, pericytes, and astrocytes [20, 31]. Once inside the brain, metastatic cells adapt to the unique neural microenvironment by leveraging local signaling, angiogenesis, and metabolic support systems [20]. Despite therapeutic advances, treating melanoma brain metastases remains difficult due to the BBB and the additional barrier imposed by the tumor microenvironment [32]. Even in cases where the BBB is compromised by the tumor, it remains uncertain which therapeutic agents can reach effective concentrations at the tumor site.

Recently, Gonzalez et al. analyzed gene expression on the single-cell level in brain metastases from established primary tumors demonstrating distinctive and shared features across metastases [33]. Karreman and Winkler provided an insightful review of the cancer neuroscience of brain metastasis, exploring how the neural system can control metastatic tumor cells that originate from outside the brain, as in the case of melanoma, and how they can plastically modify the brain [34]. Here we go down to protein molecules and signaling pathways. Given the complexity and heterogeneity of melanoma brain metastases, elucidating the genomic alterations that drive brain colonization is critical for uncovering targetable molecular drivers. Toward this crucial aim, we compare the mutational landscapes of primary melanoma tumors, metastatic melanoma, and metastatic brain tumors without lineage match to uncover brain-specific alterations and affected pathways. Our analytical strategy first identifies mutations enriched during progression from primary to metastatic stages, followed by pathway enrichment analyses to reveal dysregulated signaling networks. By integrating mutation and pathway data, we aim to infer mechanisms associated with brain colonization and define molecular features that distinguish brain metastases. By integrating and comparing datasets, here we identify pathways and molecular dependencies unique to brain metastases, including neurotrophin, MAPK, and PI3K/AKT signaling, which may inform the development of more effective combination therapies.

## Results

### Mutated proteins in primary/metastatic melanoma and metastatic brain tumors

To investigate shared and distinct mutational patterns across melanoma progression stages, we analyzed mutation frequencies in 256 primary melanoma, 474 metastatic melanoma, and 360 brain metastatic tumors (with samples likely mostly from lung, breast, and leukemia, in addition to e.g., prostate, colon, and kidney tumors, though no details are provided in available statistics, possibly due to sparsity of clinical samples). In total, we identified 644 mutated proteins in primary melanoma (2.51 mutations per tumor), 1,458 in metastatic melanoma (3.07 mutations per tumor), and 613 in brain metastases (1.80 mutations per tumor) (Figure 1, left). The values in parentheses represent the average number of mutated proteins per tumor sample within each group, reflecting an overall *increase in mutational burden from the primary melanoma tumor to its metastases, followed by a reduction in brain metastases*. When focusing on recurrently mutated proteins within each cancer type, we observed 231 in primary melanoma (0.90 per tumor), 347 in metastatic melanoma (0.73 per tumor), and 221 in brain metastases (0.61 per tumor) (Figure 1, right). *This progressive reduction in recurrently mutated proteins per tumor suggests a narrowing of the mutational repertoire with disease progression*. Such a trend may reflect clonal selection during metastasis, in which only subsets of tumor cells carrying advantageous mutations successfully colonize distant sites. In the brain, this may indicate reliance on a more restricted set of driver alterations optimized for survival in the unique microenvironment. The mutations are listed in Supplementary Data 1.

**Figure 1.**
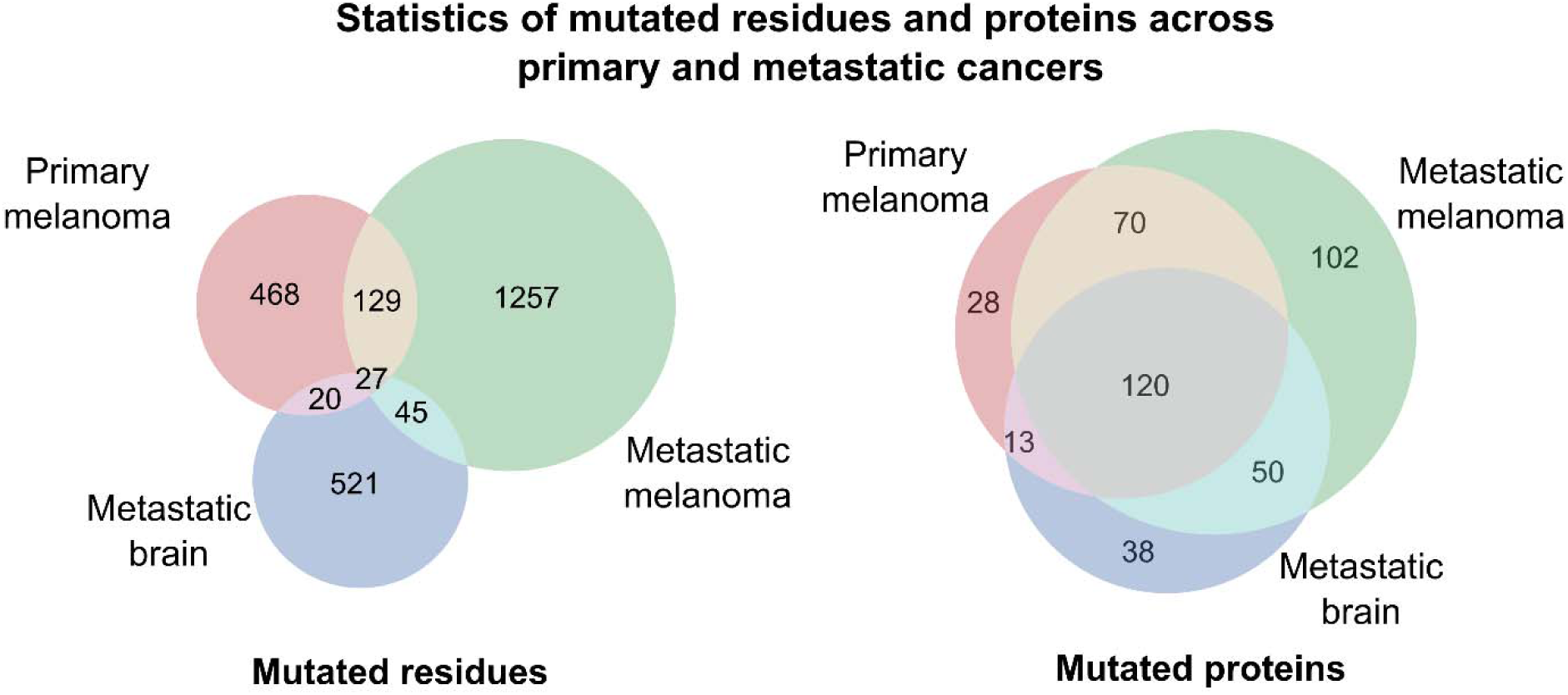
Mutation frequencies across primary melanoma, metastatic melanoma, and metastatic brain tumors. 644 mutated proteins in primary melanoma (2.51 mutations per tumor), 1,458 in extracranial metastatic melanoma (3.07 mutations per tumor), and 613 in brain metastases (1.80 mutations per tumor) (left). Focusing on proteins with mutation frequencies within each cancer type, we observed 231 mutated genes in primary melanoma tumors (0.90 mutated genes per tumor), 347 in metastatic melanoma tumors (0.73 mutated genes per tumor), and 221 in brain metastases (0.61 mutated genes per tumor) (right). This reduction in mutated proteins per tumor suggests a narrowing of the mutational repertoire during disease progression. Such a trend may reflect clonal selection in metastasis, where only subsets of tumor cells with advantageous mutations successfully colonize distant sites. In the brain, this may point to dependence on a more restricted set of driver alterations optimized for survival within the unique microenvironment.

Among the 12 proteins with relative frequencies, notable patterns emerge: isocitrate dehydrogenase 1 (*IDH1*) and *TP53* show disproportionately high mutation frequencies in brain metastases (29.72% and 31.67%, respectively), while *BRAF* and *NRAS* mutations dominate in melanoma metastases and primary melanoma tumors. *PTEN, NF1, ERBB4, CDKN2A* and *PIK3CA* exhibit more balanced frequencies across tumor types, potentially pointing to broader roles in melanoma metastatic progression with no restriction to one tumor type. Among the mutated proteins, only 120 were shared across all tumor types, and only 12 of these reached ≥4.5% mutation frequency in at least one cancer type, *highlighting the molecular divergence that accompanies metastasis, particularly to the brain* (Figure 2 presents a 3D scatter plot). Proteins with mutation frequencies below 4.5% are shown in Figure S1, and the corresponding frequencies are listed in Supplementary Data 2. Pathway enrichment analysis of the 120 proteins commonly mutated across primary melanoma, metastatic melanoma, and brain metastases using MSigDB Hallmark gene sets revealed convergence on core oncogenic and tumor suppressor pathways (Figure S2). The most significantly enriched pathways included Transcription Factor 1 (E2F1) targets, indicating dysregulated cell cycle progression and increased proliferative signaling [35–37]. The interplay between E2F1 and PI3K pathways is important in breast cancer brain metastasis. E2F1 is a member of E2F family of transcription factors which are upregulated in brain metastases. It regulates both proliferation and apoptosis. PI3K/AKT pathway suppresses E2F1’s apoptotic targets, thereby promoting tumor growth [38].

**Figure 2.**
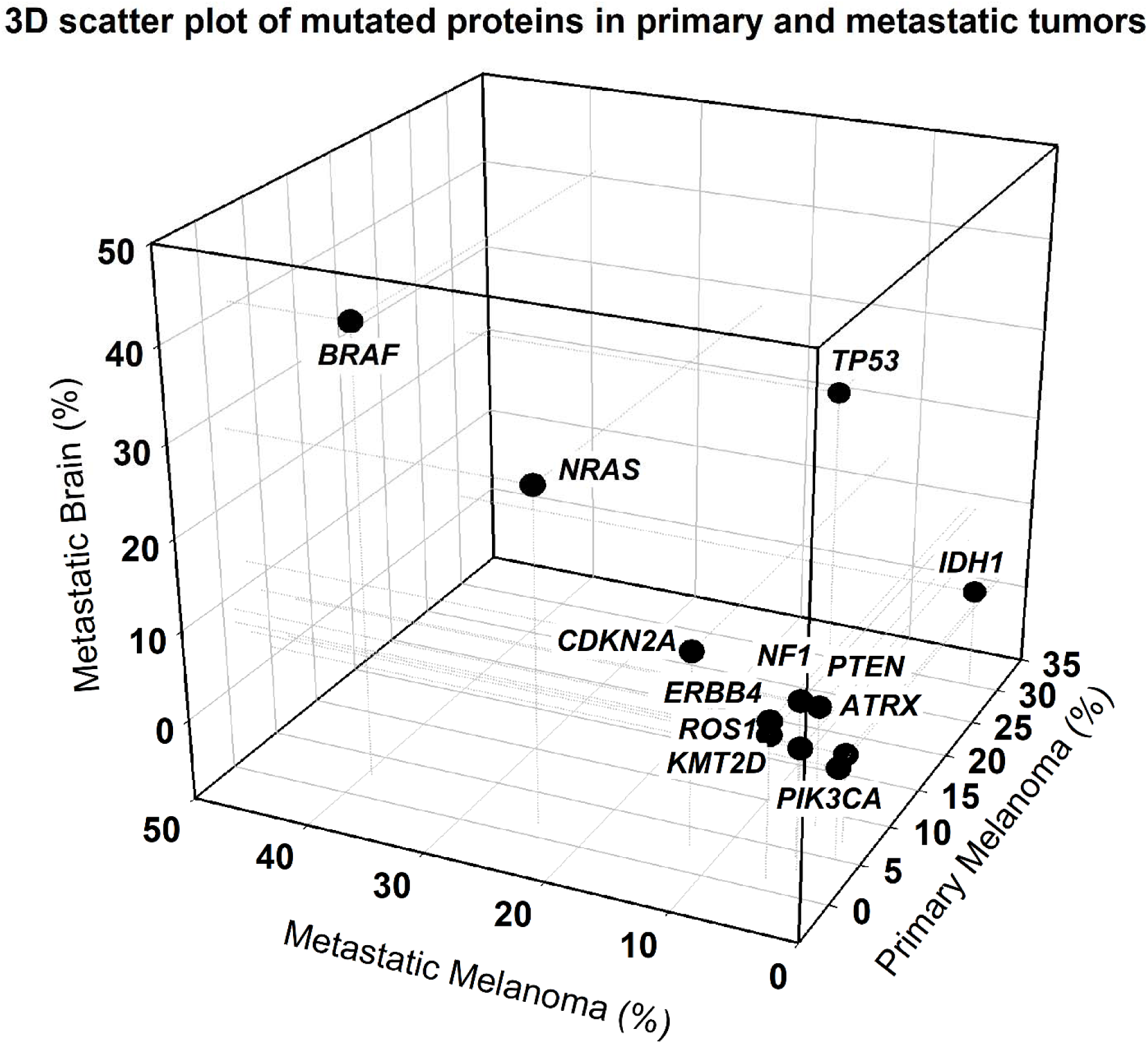
3D scatter plot of mutation frequencies. 3D scatter plot shows mutations observed in 360 brain metastatic tumor samples, 256 primary melanoma tumors, and 474 metastatic melanoma tumors. Mutation data includes 613 mutated proteins in brain metastasis tumors, 644 in primary melanoma tumors, and 1,458 in metastatic melanoma tumors. Among these, 120 mutated proteins are shared by all three cancer types where 12 out of these commonly mutated proteins have any mutation frequency ≥4.5%, including two with mutation frequencies ≥4.5% across all tumor types. Genes mutated at ≥4.5% in at least one tumor type include ATRX (6.94% in brain, 2.34% in primary, 1.05% in metastatic melanoma), BRAF (2.5%, 39.45%, 40.72%), CDKN2A (2.78%, 12.5%, 12.66%), ERBB4 (1.11%, 5.47%, 8.44%), IDH1 (29.72%, 2.34%, 2.53%), KMT2D (2.78%, 3.91%, 4.64%), NF1 (4.17%, 4.69%, 8.44%), NRAS (1.39%, 24.22%, 28.06%), PIK3CA (5.0%, 1.95%, 1.27%), PTEN (8.06%, 5.08%, 4.64%), ROS1 (1.11%, 5.47%, 6.96%), and TP53 (31.67%, 15.63%, 21.52%). The 3D scatter plot visualizes these genes’ mutation frequencies (≥4.5% in at least one cancer type) in Primary Melanoma (x-axis, light blue), Metastatic Melanoma (y-axis, dark blue), and Brain Metastases (z-axis, magenta). Each point corresponds to a protein positioned by its mutation frequency across the three cancer types. Dotted projection lines help trace each protein’s position relative to other axes, and protein names are annotated at their coordinates for clarity.

Additional enrichment was observed in Notch, Hedgehog, and Wnt/β-catenin signaling pathways, all of which play key roles in cell fate determination, stemness, and metastasis [39–41]. Enrichment of the G2-M checkpoint and p53 pathways suggests impaired genomic stability and apoptosis[42–44]. Mutations in genes associated with *KRAS* and PI3K/AKT signaling support activation of growth and survival programs [45, 46]. These findings highlight the convergence of shared mutations on critical regulatory circuits across these primary and metastatic tumor subtypes.

We examined the 120 commonly mutated genes for their presence in the Human Metastatic Database [47], which catalogs 286 metastatic genes. Five genes -*CTNNB1*, *ERBB3, FGFR1, PDGFRA,* and *PDGFRB*-were shared across three cancer types, highlighting their consistent involvement in metastatic biology. Additional overlaps included *CDH1, INHBA, PAX5, RUNX1T1* (shared between primary and metastatic melanoma tumors), *CD79A, NKX2-1* (shared between melanoma primary tumors and brain metastases), and *SLC34A2* (shared between melanoma and brain metastases). The recurrence of growth factor receptors (*ERBB3, FGFR1, PDGFRA, PDGFRB*) and the signaling mediator *CTNNB1* suggests these genes act as core drivers of metastatic dissemination and colonization in melanoma [48, 49].

Several of these proteins play critical roles in brain metastases by modulating tumor progression, therapy resistance, and the tumor microenvironment. *CTNNB1* facilitates BBB penetration and contributes to immune evasion and therapy resistance via Wnt/β-catenin, epithelial–mesenchymal transition (EMT), and PI3K/AKT signaling [50–52]. *ERBB3* promotes survival and therapy resistance through PI3K/AKT, MAPK, and other resistance pathways [53]. *FGFR1* enhances proliferation and MAPK inhibitor resistance, increasing brain metastatic potential via FGF/FGFR, MAPK, and VEGF signaling [54–56]. *PDGFRA* supports angiogenesis, BBB penetration, and vascular remodeling through PDGF/PDGFR, VEGF, and tumor-stroma interactions [57–59].

Subsequently, we investigated mutations in the most recurrently altered proteins. Analysis of *TP53* across melanoma progression stages uncovered shared as well as distinct mutational features (Figure 3a). Primary melanomas harbored recurrent mutations at H179 and R213, while melanoma metastases accumulated a broader spectrum of mutations, including R342, R248, S127, C242, P278, R110, R196, and S241. Brain metastases showed even greater heterogeneity, with mutations spanning residues H179, R213, R342, R248, R241, R175, V173, G245, R273, P152, and Y234, suggesting that *selective pressures in the brain microenvironment may favor the expansion of TP53-mutant subclones*. Mapping these variants onto the *TP53* protein highlights their distribution across key functional domains (Figure 3b). Mutations within the DNA-binding domain (e.g., R175, R213, R248, R273, G245) compromise *TP53*’s ability to regulate DNA repair, apoptosis, and cell-cycle checkpoints, while alterations in the oligomerization domain (e.g., R342) may destabilize tetramer formation and functional activity [60, 61]. Collectively, these patterns indicate that *TP53* loss of function is not only a recurrent event in melanoma but also evolves with disease progression, with brain metastases exhibiting particularly disruptive variants. This suggests that TP53 inactivation may facilitate both early tumor initiation and metastatic adaptation, where disruption of genomic surveillance, apoptotic signaling, and DNA repair pathways could provide the selective advantage for tumor cells to survive in hostile metastatic niches, including the brain. This raises the possibility that *TP53*-mutant melanomas may exhibit heightened resistance but also harbor vulnerabilities to targeting defective DNA repair machinery.

**Figure 3.**
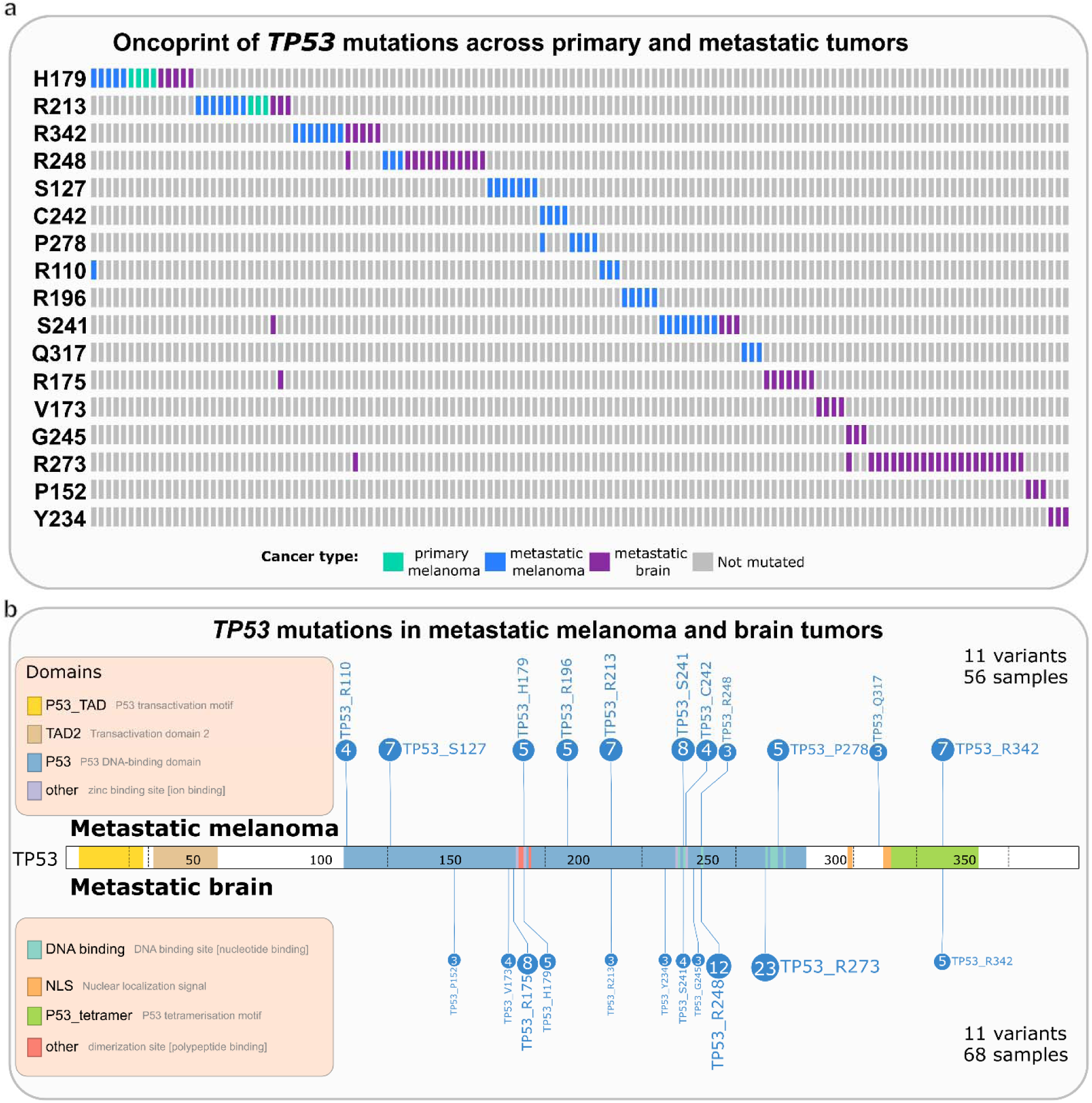
*TP53* mutations in primary melanoma, metastatic melanoma and metastatic brain tumors. **(a)** In the oncoprint, each bar in columns represents a tumor harboring the corresponding mutation in primary melanoma (green), metastatic melanoma (blue), metastatic brain (purple), gray bars indicate no mutation. Metastatic melanoma harbors mutations in H179, R213, R342, R248, S127, C242, P278, R110, R196, S241. Primary melanoma has mutations H179, R213 and metastatic brain tumors have mutations in H179, R213, R342, R248, S241, R175, V173, G245, R273, P152, Y234. **(b)** TP53 mutation distribution in metastatic melanoma and brain metastases. Lolipop plot shows TP53 mutations occurring at residues R342, H179, R248, R213, S127, C242, P278, R110, R196, S241, and Q317, spanning key functional domains of the protein, including the transactivation domain, DNA-binding domain, and oligomerization domain. Mutations in the DNA-binding domain impair *TP53*’s regulation of DNA repair, apoptosis, and cell-cycle checkpoints, while alterations in the oligomerization domain may disrupt tetramer formation and functional activity. Lollipop plot figure id prepared using the ProteinPaint tool (https://proteinpaint.stjude.org).

A random forest classifier trained on binary mutation profiles demonstrated the highest predictive performance for metastatic melanoma (precision = 0.52, recall = 0.91), followed by brain metastases (precision = 0.89, recall = 0.43), with poor classification of primary melanoma (precision = 0.24, recall = 0.06), yielding an overall accuracy of 56% (n = 255) (Figure S3). Feature importance analysis revealed distinct sets of mutations driving classification across tumor types. Mutations in *BRAF*, *NRAS*, and *NF1* were most informative for metastatic melanoma, reflecting their role as dominant oncogenic drivers in progression. Brain metastases were strongly influenced by mutations in *CTNNB1*, *ERBB3*, *FGFR1*, *PDGFRA*, and *PDGFRB*, genes implicated in BBB penetration, survival in the neural microenvironment, and metastatic colonization [62, 63]. In contrast, classification of primary tumors relied more on *CDKN2A*, *TP53*, and *PTEN*, consistent with early disruption of tumor suppressor pathways during melanoma initiation [9, 64]. These results suggest that while overlapping mutations blur distinctions between tumor stages, subsets of driver mutations provide discriminatory power that reflects their roles: proliferation and signaling activation in melanoma metastases, microenvironmental adaptation in brain metastases, and tumor suppressor loss in primary melanoma tumors.

Analysis of recurrent oncogenic drivers revealed distinct mutation patterns across melanoma progression. In metastatic melanoma, *BRAF* and *TP53* mutations are mutually exclusive (Figure S4). *BRAF*^V600^ represented the dominant activating alteration, while non-V600 variants at residues D594, G466, G469, L597/K601, and S467 highlighted the functional heterogeneity of *BRAF* mutations, spanning strongly activating, constitutively dimerizing, and kinase-impaired classes with different signaling dependencies [65, 66]. The mutual exclusivity of the aggressive *BRAF* and *TP53* alterations likely reflects distinct evolutionary trajectories: tumors either rely on MAPK pathway hyperactivation through BRAF mutation or tolerate genomic instability through TP53 loss, but rarely both [67, 68]. By contrast, *KRAS* mutations (G12 and Q61) were observed in both primary tumors and brain metastases, with Q61 predominating in primary tumors and the generally more aggressive G12 variants distributed across primary melanoma and metastatic brain tumors (Figure S5). In brain metastases, *IDH1* and *TP53* mutations frequently co-occurred (Figure S6), particularly with canonical, weaker *IDH1*^R132^ alterations, which have better prognosis. Mapping *BRAF* and *NRAS* mutations across tumor types underscored distinct trends: *BRAF*^V600^ is the most common alteration across three cancer types, while non-V600 variants were enriched in metastatic tumors (Figure S7). Similarly, *NRAS*^Q61^ was the dominant mutation across all tumor types, whereas *NRAS*^G12^ and *NRAS*^G13^ were less frequent and more often associated with metastases (Figure S8). Co-occurrence of highly aggressive mutations can promote OIS (oncogene induced senescence), thus rarer [69].

Collectively, these patterns indicate that melanoma metastasis can arise through distinct oncogenic routes such as *BRAF*-driven MAPK hyperactivation, *TP53*-driven genomic instability, or cooperative alterations like *IDH1*–*TP53* in brain metastases. *These observations suggest that melanoma progression is propelled by mutations converging on a restricted set of regulatory circuits*, enabling coupling uncontrolled proliferation with increased plasticity and survival. The recurrent involvement of signaling receptors including *ERBB3*, *FGFR1*, *PDGFRA*, and *PDGFRB*, together with *CTNNB1* suggests that targeting shared signaling vulnerabilities, rather than site-specific alterations, could provide therapeutic opportunities to disrupt metastatic progression across multiple tumor contexts.

### Dissection of signaling mechanisms in melanoma and brain metastasis through mutated pathways

After dissecting mutated residues in several key proteins across primary melanoma, metastatic melanoma, and metastatic brain tumors, we mapped these proteins to signaling pathways. Using the KEGG 2021 pathway dataset [70], we identified signaling pathways with elevated mutation load in the three tumor types. Figure 4 shows a 3D scatter plot depicting the mutation frequencies of signaling pathways across these cancers (Supplementary Data 2). In total, 18 pathways are commonly mutated in all three cancer types. Neurotrophin signaling showed the highest mutation frequency in primary (76.6%) and metastatic melanoma (79.5%) but was lower in brain metastases (43.1%). Rap1 and ErbB signaling were also highly mutated in melanoma (74.2–77.4% and 73.8–75.7%, respectively) but much less so in brain metastases (18.3–20%). MAPK signaling remained frequently mutated across all types (66.4% in primary, 70.9% in metastatic melanoma, and 47.8% in brain metastases). PI3K/AKT signaling showed relatively consistent mutation frequencies (55.1% in primary, 59.5% in metastatic melanoma, and 51.1% in brain metastases). FoxO and Phospholipase D pathways were moderately mutated in melanoma (51.6–52.5% and 37.1–40.9%, respectively) but reduced in brain metastases (19.4% and 18.9%). cAMP, mTOR, and Relaxin signaling were also less frequently mutated in brain metastases (13.1–14.4%) compared to melanoma (35.2–50%). Prolactin and AGE-RAGE signaling showed modest mutation in melanoma (32.4–36.5%) but declined in brain metastases (11.1–11.9%). In contrast, p53 and thyroid hormone pathways exhibited relatively higher mutation load in brain metastases (41.4% and 37.2%) compared to melanoma (31.6–39.7% and 25.4–32.1%, respectively). Wnt and sphingolipid signaling also showed increased mutations in brain metastases (35.3–35.6%) relative to melanoma (22.7–29.3%). JAK/STAT and HIF-1 signaling remained low across all tumor types (12.9–19.4%). This analysis highlights both conserved and divergent pathway-level alterations during melanoma progression and metastasis.

**Figure 4:**
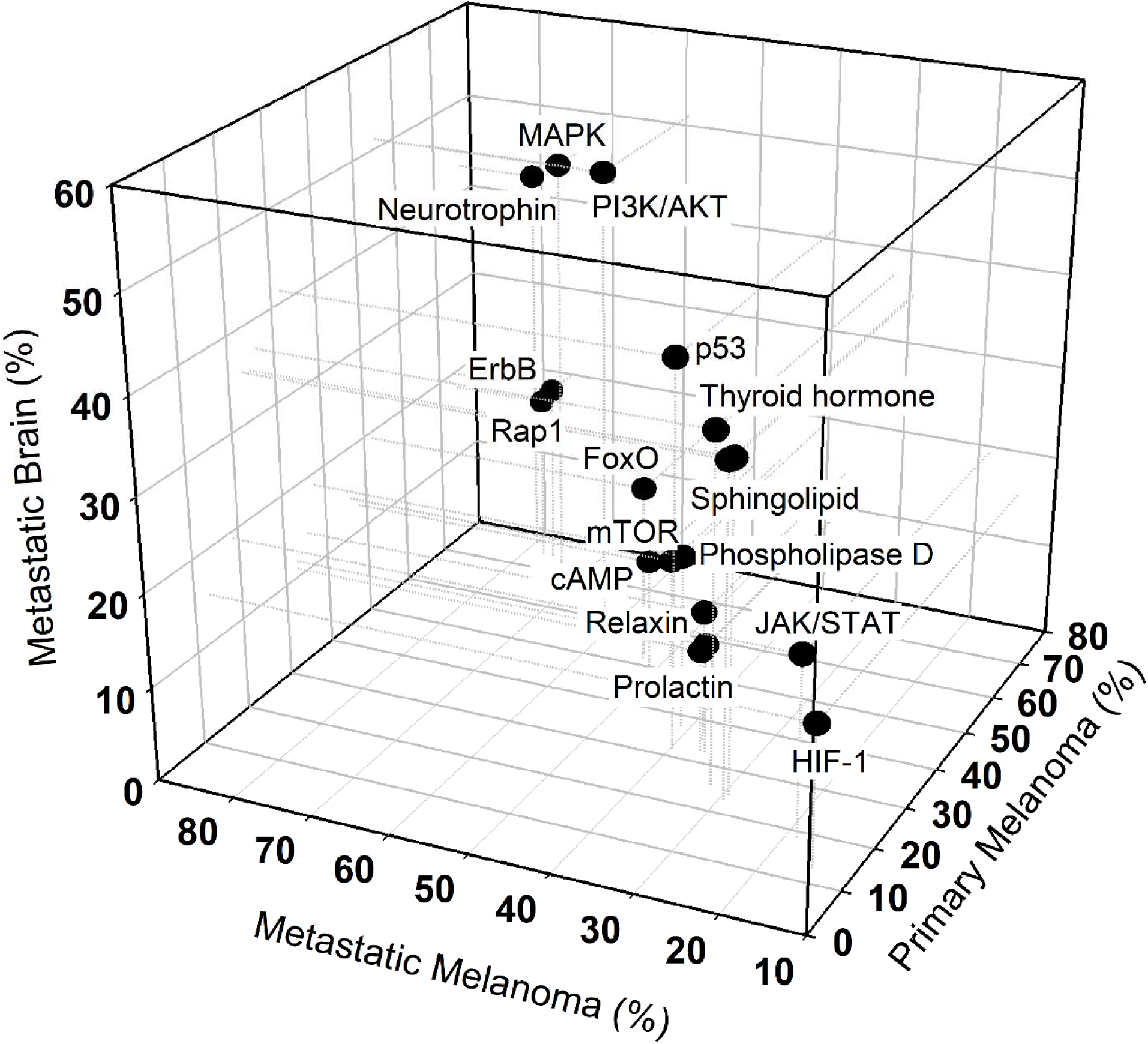
3D scatter plot of pathway mutations in primary melanoma, metastatic melanoma and brain tumors. Each dot represents a signaling pathway commonly mutated across the three cancer types. The x-, y-, and z-axes show the percentage of tumors with mutations in pathway proteins in primary melanoma, metastatic melanoma, and brain metastases, respectively. Dotted projection lines aid spatial interpretation, and pathway names are annotated in 3D. Pathways with mutation frequency >0% in at least one condition are included, totaling 22. Rap1, MAPK, and Neurotrophin signaling pathways were highly mutated in primary and metastatic melanoma (≥76%) but less so in brain metastases (24–42%). ErbB, FoxO, and Insulin signaling showed similarly high mutation in melanoma (72–75%) but lower rates in brain metastases (16–24%). PI3K/AKT and Thyroid hormone signaling were consistently mutated across all three tumor types (≥49% in brain). Wnt, Sphingolipid, and p53 signaling showed moderate mutation in primary melanoma (24–32%) but higher levels in brain metastases (35–41%). In contrast, cAMP, Estrogen, and Relaxin signaling were less frequent in brain metastases (10–14%) despite being more common in melanoma. JAK/STAT signaling remained consistently low across all types (<16%).

The complexity of cancer therapy appears in part from interconnected signaling pathways in cancer and innate and adaptive immunity [71, 72]. Neurotrophins are a family of growth factors whose receptors, especially Trk receptors and p75^NTR^, support cancer cell survival, proliferation, invasion, and treatment resistance [73]. Specific neurotrophins, such as nerve growth factor (NGF) and NT-3, promote invasion, with melanoma cells expressing receptors like p75^NTR^ and TrkC mediating these effects [73]. Metastatic melanoma cells exhibit increased proliferation when NGF activates the MAPK pathway via TrkA [74, 75]. In brain-metastatic melanoma cells, neurotrophins stimulate the production of extracellular matrix (ECM)–degrading enzymes. These cells also secrete autocrine and paracrine factors that regulate their own growth and survival, as well as neurotrophin activity in surrounding brain cells, including astrocytes [76]. Elevated neurotrophin in melanoma cells may facilitate brain invasion [73, 77, 78]. Neurotrophins also help brain microenvironment homeostasis; their receptors TrkB and TrkC act as proto-oncogenes, and disruptions in their signaling can affect apoptosis [79].

We next illustrate selected signaling pathways (Figures 5–6), highlighting cancer type–specific and shared mutations, with a focus on PI3K/AKT, MAPK, and JAK/STAT pathways. These are presented in connection with neurotrophin signaling in melanoma brain metastasis, particularly invasion and angiogenesis. In primary melanoma (Figure 5, top panel), recurrent mutations cluster in the JAK/STAT and MAPK pathways, underscoring their central role in tumor initiation and progression [41, 80]. Alterations in the receptor tyrosine kinase c-KIT and in VEGF/VEGFR signaling components initiate aberrant signaling, which is amplified by downstream mutations in Ras and *BRAF [58, 81–83]*. The *BRAF*^V600^ variants, along with other activating BRAF mutations, represent key oncogenic drivers of melanoma. *BRAF*^V600^ is observed in three cancer types while other weaker non-V600 variants (D594, G466, G469, L597, S467) are less common in metastatic tumors. *VEGF/VEGFR* signaling engages the PI3K/AKT cascade, resulting in crosstalk between MAPK and PI3K/AKT pathways that converge on transcriptional regulators such as STAT proteins and c-MYC [58, 84]. These alterations foster uncontrolled proliferation, survival, and resistance to apoptosis in primary tumors, while also conducive to metastatic dissemination. In metastatic brain tumor samples (Figure 5, bottom panel), mutations in *EGFR* and *TSC2* suggest dysregulation of growth factor signaling and mTOR pathways.

**Figure 5.**
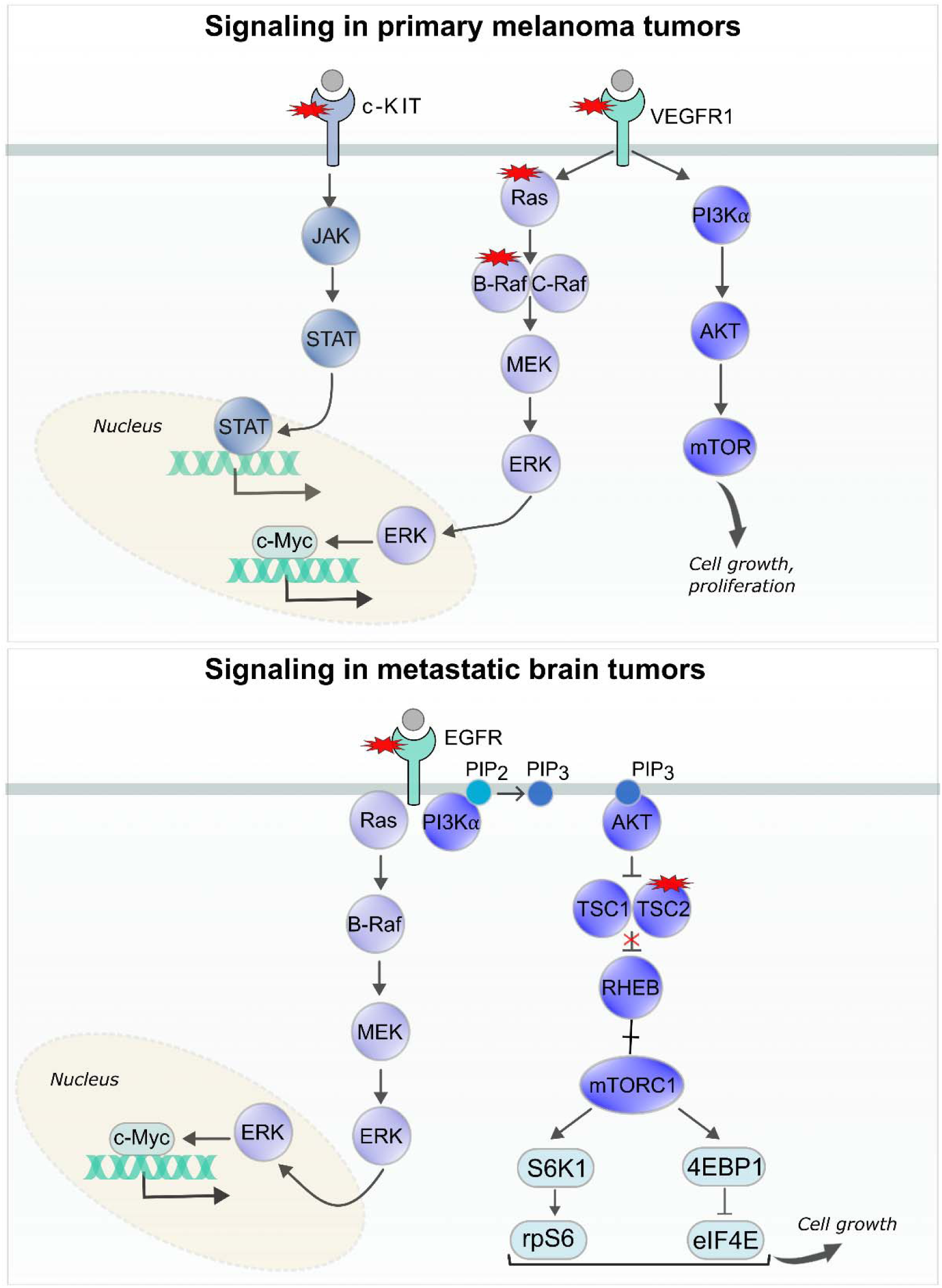
Mutational landscape of signaling pathways in primary melanoma and brain metastases. **(Top)** In the JAK/STAT signaling pathway, the receptor c-KIT is mutated in primary melanoma samples (top panel). Within the MAPK pathway, mutations are observed in *VEGF* and its receptor *VEGFR*. Downstream components, including Ras and *BRAF*, are also mutated; specific BRAF mutations include V600E, V600K, and other activating variants frequently observed in melanoma. Signaling downstream of *VEGF* and *VEGFR1* activates both the MAPK and PI3K/AKT pathways. These cascades converge on transcriptional regulators such as STAT proteins and c-MYC, driving proliferation and survival. **(Bottom)** In metastatic brain tumors (bottom panel), mutations in *EGFR* and *TSC2* suggest disrupted growth factor signaling and dysregulated mTOR activity. Both primary and metastatic tumors share mutations in p14^ARF^ and p53, key regulators of cell cycle control and apoptosis. Additional shared mutations in *BRAF*, *PTEN*, and p16^INK4A^ (*CDKN2A*) indicate disruption of multiple tumor suppressor pathways. Color coding denotes major signaling pathways: dark purple, PI3K/AKT pathway; light purple, MAPK pathway; light blue, JAK/STAT pathway. Red star symbols mark mutated proteins within these pathways.

**Figure 6.**
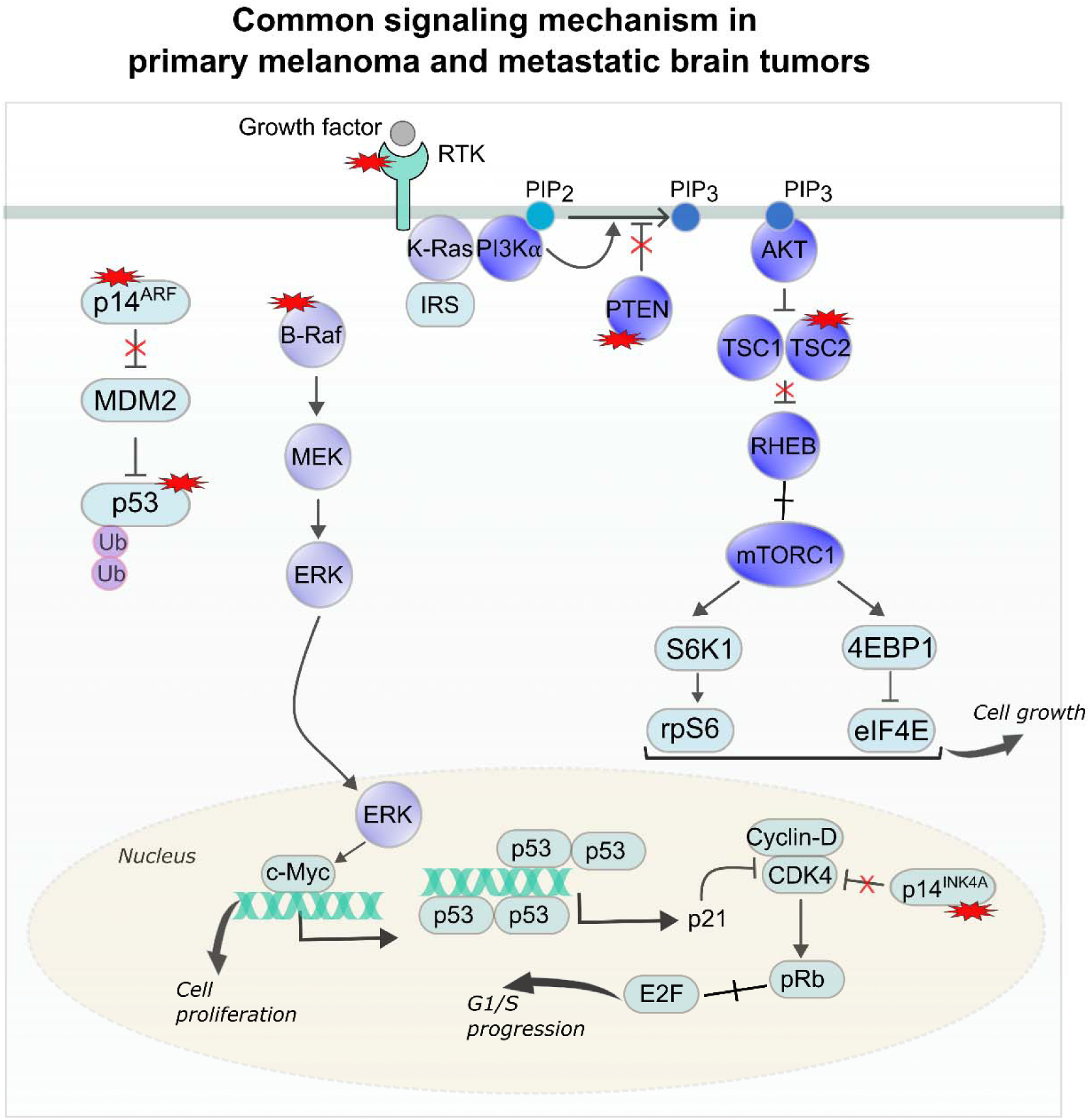
Brain metastases harbor mutations in growth factor signaling genes such as *EGFR* and *TSC2,* affecting the mTOR axis. Shared tumor suppressor mutations include p14^ARF^, p53, and p16^INK4A^, along with common oncogenic drivers *BRAF*^V600^ and *PTEN*. MDM2-mediated *TP53* degradation may contribute to the elevated p53 mutation burden. Alterations in PI3K/AKT and MAPK pathways promote proliferation, survival, and therapy resistance, while p53 loss impairs DNA repair, apoptosis, and cell-cycle control, fostering genomic instability. Neurotrophin signaling via Trk receptors (NTRK1/2/3) and p75^NTR^ recruits adaptor proteins (Shc, Grb2) to activate Ras/MAPK and PI3K/AKT, enhancing proliferation, differentiation, survival, and anti-apoptotic signaling. Potential therapeutic strategies include co-targeting c-KIT and VEGFR to inhibit growth-promoting pathways. Color coding denotes major signaling pathways: dark purple, PI3K/AKT pathway; light purple, MAPK pathway; light blue, p53 pathway. Red star symbols mark mutated proteins within these pathways.

In melanoma brain metastases (Figure 6), the mutational spectrum shifts toward growth factor signaling pathways that enhance adaptation to the brain microenvironment. Recurrent mutations in *EGFR* and *TSC2* implicate dysregulation of the mTOR axis, while additional alterations in PI3K/AKT and MAPK pathways reinforce drug-resistance and pro-survival [85–87]. Shared disruptions in tumor suppressors, including p14^ARF^, p53, and p16^INK4A^ (encoded by *CDKN2A*), alongside drivers such as *BRAF*^V600^ (class I) and *PTEN* mutations, highlight vulnerabilities in primary melanoma and metastatic disease [88, 89]. *MDM2*-mediated *TP53* degradation may amplify p53 dysfunction, genomic instability and impaired DNA repair [90–92]. Brain metastases also engage neurotrophin signaling via Trk receptors (NTRK1/2/3) and p75^NTR^, which activate Ras/MAPK and PI3K/AKT cascades to promote survival and anti-apoptotic signaling [73, 77].

In primary melanoma, *BRAF*^V600E^ mutations were observed in 66 tumors, whereas *BRAF*^V600K^ mutations were detected in 13 tumors. In metastatic melanoma, the distribution was broader: 125 tumors harbored V600E, 22 tumors V600K, 3 tumors V600M, and 5 tumors V600R. Similarly, in metastatic *BRAF*-driven tumors, V600E predominated with 6 cases. B-Raf is frequently mutated in melanoma, with V600E occurring more commonly (60%-80%) than the more aggressive V600K (10%-30%) due to a single versus double nucleotide change, respectively [93, 94]. Although both mutations occur at the same kinase domain residue and were initially thought to behave similarly, emerging data show that V600K tumors are more aggressive, have higher relapse risk, shorter survival, and differ in drug responsiveness compared to V600E tumors [95]. Both *BRAF* V600E and V600K mutants can be constitutively active as monomers, but dimerization enhances MAPK signaling. Mutations at V600 disrupt the N-lobe hydrophobic surface in the inactive OFF-state and favor the active ON-state without destabilizing residue contacts, with V600E forming stabilizing salt bridges absent in V600K [95], suggesting that subtle structural differences may underlie distinct clinical behaviors.

These highlight therapeutics in advanced melanoma beyond single-target inhibition, targeting converging pathway combinations, such as combining c-KIT and *VEGFR* inhibitors. Neurotrophin receptors, along with MAPK and PI3K/AKT pathway components, also emerge as key targets in disease progression. The PI3K/AKT pathway promotes melanoma, and its activity is higher in the brain microenvironment than elsewhere[96]. Preclinical models demonstrate that inhibition of this pathway reduces tumor growth, induces apoptosis, and when combined with *BRAF* inhibition, significantly prolongs survival in mice with brain metastases. Co-targeting neurotrophin receptors and MAPK pathway members, particularly with *BRAF* inhibitors, represents a promising combination strategy for melanoma brain metastases. Overall, these underscore the molecular heterogeneity of melanoma brain metastases, driven by interconnected signaling and reveal targets that may differ from those in other melanoma metastases.

## Discussion

We aim to elucidate the mechanistic underpinnings of melanoma brain metastasis through mutations in cellular signaling pathways. We identified distinct pathway-level alterations across melanoma progression: Rap1 and ErbB signaling are highly mutated in primary and metastatic melanoma but diminished in brain metastases, whereas p53, thyroid hormone, Wnt, and sphingolipid signaling became more prominent. In contrast, neurotrophin, MAPK, and PI3K/AKT signaling are consistently prominent across all three cancer types.

Mechanistically, neurotrophin signaling emerged as a central driver: neurotrophins and their receptors, especially Trk and p75^NTR^, regulate melanoma survival, invasion, and resistance [73], with brain-metastatic cells exploiting these cues to upregulate the extracellular matrix and engage astrocytic support [76, 77]. Thyroid hormone signaling further promotes proliferation, angiogenesis, and epithelial–mesenchymal transition [97–100], linking endocrine regulation to metastasis, while also intersecting with neurotrophin pathways that shape brain development [101, 102]. Metastatic melanoma, particularly brain metastases, showed a broader mutational spectrum and enrichment of PI3K/AKT activity, supporting survival and adaptation to the brain microenvironment [16, 103]. Recurrent alterations in *ERBB3*, *FGFR1*, and *PDGFR*s indicate that melanoma progression converges on shared signaling hubs despite diverse oncogenic routes.

### Therapeutic Potential of PI3K/AKT Inhibition and MAPK Pathway Resistance Dynamics

In vivo imaging revealed PI3K/AKT/mTOR activation during early brain colonization, supporting intravascular arrest and extravasation [104]. This pathway is selectively hyperactivated in brain metastases, partly via astrocyte-derived signals [15]. Inhibition with buparlisib reduced *AKT* activity, proliferation, and tumor growth, while inducing apoptosis in mouse models, independent of *BRAF* and *NRAS* mutational status. In melanomas driven by the *BRAF*^V600E^ mutation and loss of Cdkn2a, activated *AKT1* induced lung and brain metastases, while additional loss of *PTEN* further accelerated tumor progression [105]. Both act through distinct mechanisms, underscoring PI3K/AKT as a central therapeutic target. Ras/MAPK pathway drives melanoma progression, therapy resistance, and CNS colonization [106]. *BRA*F^V600^ and *NRAS*^Q61^ mutations constitutively activate MAPK signaling, while in melanoma brain metastases, PI3K/AKT hyperactivation (often via PTEN loss) promotes resistance to *BRAF*-targeted therapy. Recurrent PI3K/AKT alterations within the BRAF-mutant subgroup suggest that *AKT* activation reduces the efficacy of BRAF + MEK inhibition [107].

### Neurotrophins and Microenvironmental Cues

Brain metastases remain a major therapeutic challenge, shaped not only by mutations but also by epigenetic and transcriptional adaptations to the brain microenvironment [16, 108]. Increasing evidence indicates that metastatic progression depends less on mutations and more on adaptive programs enabling survival in the CNS [108]. Melanoma cells exploit neurotrophins (NGF, BDNF, NT-3, NT-4/5) and their receptors to promote proliferation, invasion, and brain adaptation [79]. Trk receptors activate MAPK signaling, while p75^NTR^ engages NF-κB, enhancing ECM degradation and invasion. Astrocytes further reprogram melanoma cells toward a neural-like phenotype, supporting colonization [79].

Recent insights highlight how NGF–TrkA and EGF–EGFR signaling differentially govern melanoma cell fate via MAPK dynamics. Both converge on Ras/Raf/MEK/ERK, yet sustained ERK activation by NGF–TrkA promotes differentiation, whereas transient, high-intensity ERK activation by EGF–EGFR drives proliferation [109]. Tumoral heterogeneity often reflects cycles of dedifferentiation and re-differentiation (EMT–MET), enabling dissemination and colonization [3, 110]. While dedifferentiated subpopulations facilitate intravasation, metastases frequently reacquire epithelial-like traits, consistent with mesenchymal–epithelial transitions. In melanoma, these dynamic NGF–EGF–ERK programs may couple with EMT–MET cycles to sustain stem cell–like features, dormancy, and therapy resistance, reinforcing the need for therapies that disrupt phenotypic plasticity in addition to targeting MAPK and PI3K signaling [111].

### Conclusions

Preclinical studies demonstrate that combining PI3K or *AKT* inhibitors with *BRAF* inhibitors reduces tumor growth, induces apoptosis, and prolongs survival [96, 112]. Additional strategies, such as dual inhibition of *AKT* and *WEE1*, synergistically activate p53, enhance immunogenicity, and promote NK cell–mediated cytotoxicity. Likewise, co-targeting neurotrophin receptors and MAPK pathway components, including *BRAF*, represent a promising avenue.

Here we approached the melanoma metastases challenge through comprehensive analyses of its recurring mutations, mapped on its pathways. *Our innovative approach provides the landscape of aggressive melanoma*, uncovering its key mutations and pathways providing insight into melanoma’s mechanisms within its brain microenvironment, thereby assisting pharmacological strategies. Our work distinguishes itself by not only pointing to specific proteins and pathways to target, but to their more promising combinations–and suggests likely candidates [113]. Our findings, integrated with earlier clinical observations, highlight combination therapies in melanoma that integrate inhibitors of MAPK, PI3K/AKT, and neurotrophin signaling, potentially in conjunction with immunotherapy. They complement observations demonstrating distinctive and shared features across metastases [33]. They benefit from insight into the cancer neuroscience of brain metastasis, especially as to how the neural system can control invading metastatic tumor cells from outside the brain, as in melanoma, and how they can plastically modify the brain [34]. Collectively, these provide deeper understanding of cellular mechanisms underlying melanoma brain metastasis and multimodal approaches that may address both tumor-intrinsic alterations and microenvironmental adaptations. A next step could consider other common brain metastases, such as breast and lung cancers, which may share commonalities.

## Methods

We analyzed a dataset comprising 256 primary melanoma tumors, 474 extracranial metastatic melanoma samples, and 360 brain metastases obtained from the TCGA and AACR GENIE datasets. Although lineage-matched samples for the metastatic tumors were not available, this cohort provides meaningful insights into the molecular underpinnings of melanoma progression and dissemination to the brain.

For mutation profiling, we quantified the number of mutated residues within proteins as well as the total number of mutated proteins in primary melanoma, extracranial metastases, and brain metastases. Mutated proteins were then mapped onto KEGG signaling pathways [70], and 3D scatter plots were constructed to visualize their distribution. The frequency of mutations was calculated as the percentage of mutated tumors within each cancer type. For pathway enrichment analysis, we used MSigDB to analyze a set of 120 genes that were commonly mutated across the three tumor groups.

### Mutation feature selection

To identify genetic features distinguishing primary melanoma, metastatic melanoma, and brain metastases, we trained a random forest classifier using binary gene mutation profiles. Samples (n = 255) were split into training and held-out test sets, and classification performance was evaluated using precision, recall, F1-score, and overall accuracy. Feature importance was calculated using the Gini index (mean decrease in impurity), which reflects the contribution of each gene to reducing classification uncertainty across the ensemble of trees. The top 25 genes ranked by Gini importance were extracted, representing the features with the highest discriminative power in classifying tumor types. Importantly, these values indicate relative feature ranking rather than directionality of effect, thereby highlighting mutational drivers most relevant for tumor classification.

### Mapping mutations to signaling pathways

To illustrate signaling mechanisms implicated in melanoma progression and brain metastasis, we mapped mutated proteins onto canonical pathways curated from the KEGG database and scientific literature [70]. This approach enabled visualization of how recurrent mutations converge on core signaling cascades that drive tumor proliferation, survival, and adaptation to the brain microenvironment.

## Data and Code Availability

Somatic missense mutation profiles were obtained from The Cancer Genome Atlas (TCGA) [(https://gdc.cancer.gov/about-data/publications/mc3-2017)] and the AACR Project GENIE (Genomics Evidence Neoplasia Information Exchange). For signaling pathway analysis, the KEGG_2019_Human dataset was retrieved from the Enrichr library [(https://maayanlab.cloud/Enrichr/)], which includes 46 pathways categorized as signaling pathways.

## Acknowledgments

This Research was supported by the Cancer Innovation Laboratory, Center for Cancer Research, National Cancer Institute, National Institutes of Health Intramural Research Program project numbers, ZIA BC 010441 and ZIA BC 010442, and federal funds from the National Cancer Institute, National Institutes of Health, under contract HHSN261201500003I. The contributions of the NIH authors were made as part of their official duties as NIH federal employees, are in compliance with agency policy requirements, and are considered Works of the United States Government. However, the findings and conclusions presented in this paper are those of the authors and do not necessarily reflect the views of the NIH or the U.S. Department of Health and Human Services.

## Author contributions

B.R.Y., H.J., and R.N. conceived and designed the study. B.R.Y. did the data curation and visualization, analyzed the results and drafted the manuscript. B.R.Y., H.J., and R.N. validated, reviewed, and edited the manuscript. R.N. supervised the project.

