## Supplementary Figures for "Cellular Mechanisms Underlying Melanoma Brain Metastasis"

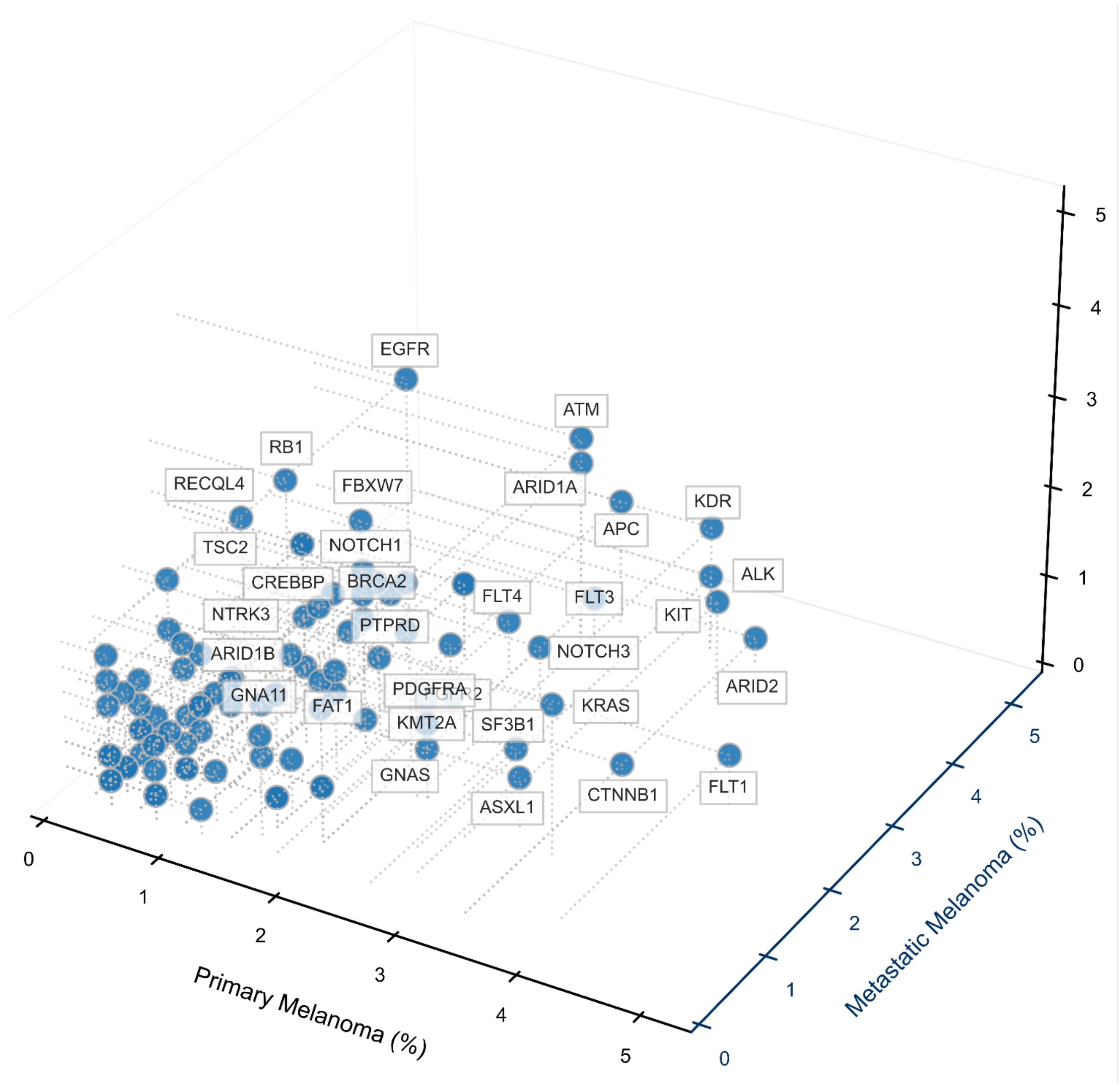


**Figure S1: 3D scatter plot of proteins with mutation frequencies below 4.5% across primary melanoma, metastatic melanoma, and metastatic brain tumors**. 3D scatter plot visualizes genes whose mutation frequencies are below 4.5% in all three tumor types: Primary Melanoma (x-axis, light blue), Metastatic Melanoma (y-axis, dark blue), and Metastatic Brain (z-axis, magenta). Each point represents a gene positioned according to its mutation percentages across the three cancer types. Presence defined as mutation frequency > 0%. Protein names are annotated when any mutation frequency exceeds 2%, enhancing interpretability.

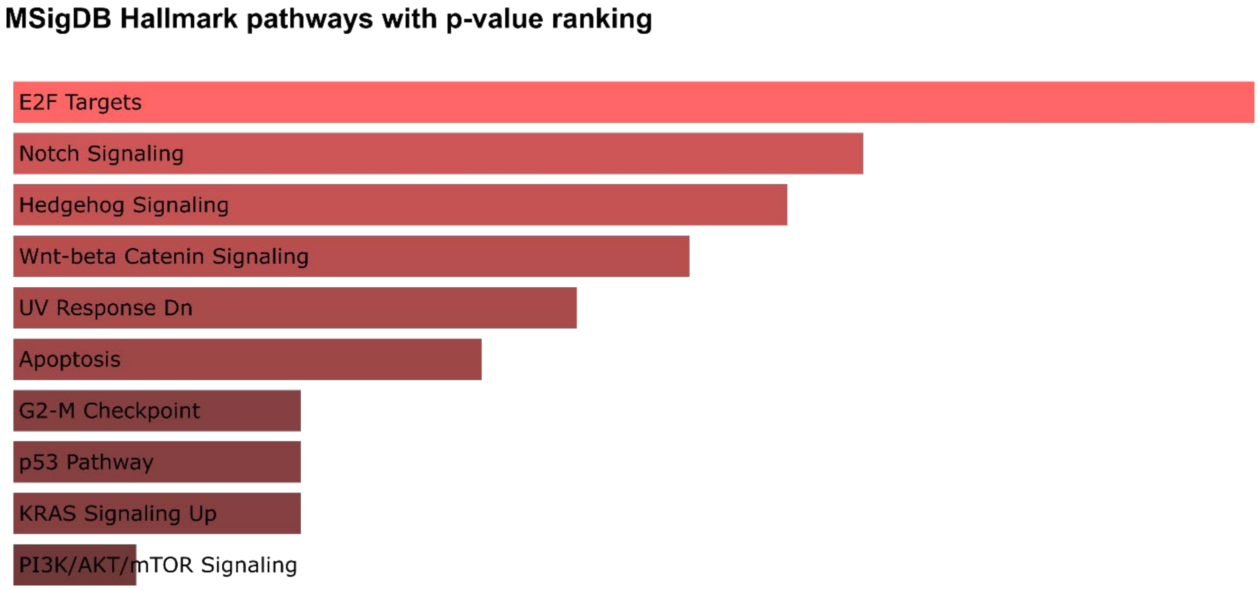


**Figure S2. Pathway enrichment analysis of 120 genes commonly mutated across samples, highlighting MSigDB Hallmark pathways ranked by statistical significance.** The analysis revealed significant enrichment of key oncogenic and tumor suppressor pathways. The 120 shared mutated genes are strongly associated with E2F targets, Notch, Hedgehog, and Wnt/β-catenin signaling, which regulate proliferation and differentiation. Pathways controlling the cell cycle and genomic stability—such as apoptosis, the G2–M checkpoint, and p53—are also enriched.

Oncogenic pathways including *KRAS* and PI3K/AKT are similarly involved, indicating that these recurrent mutations converge on core networks driving tumor progression and therapy resistance.

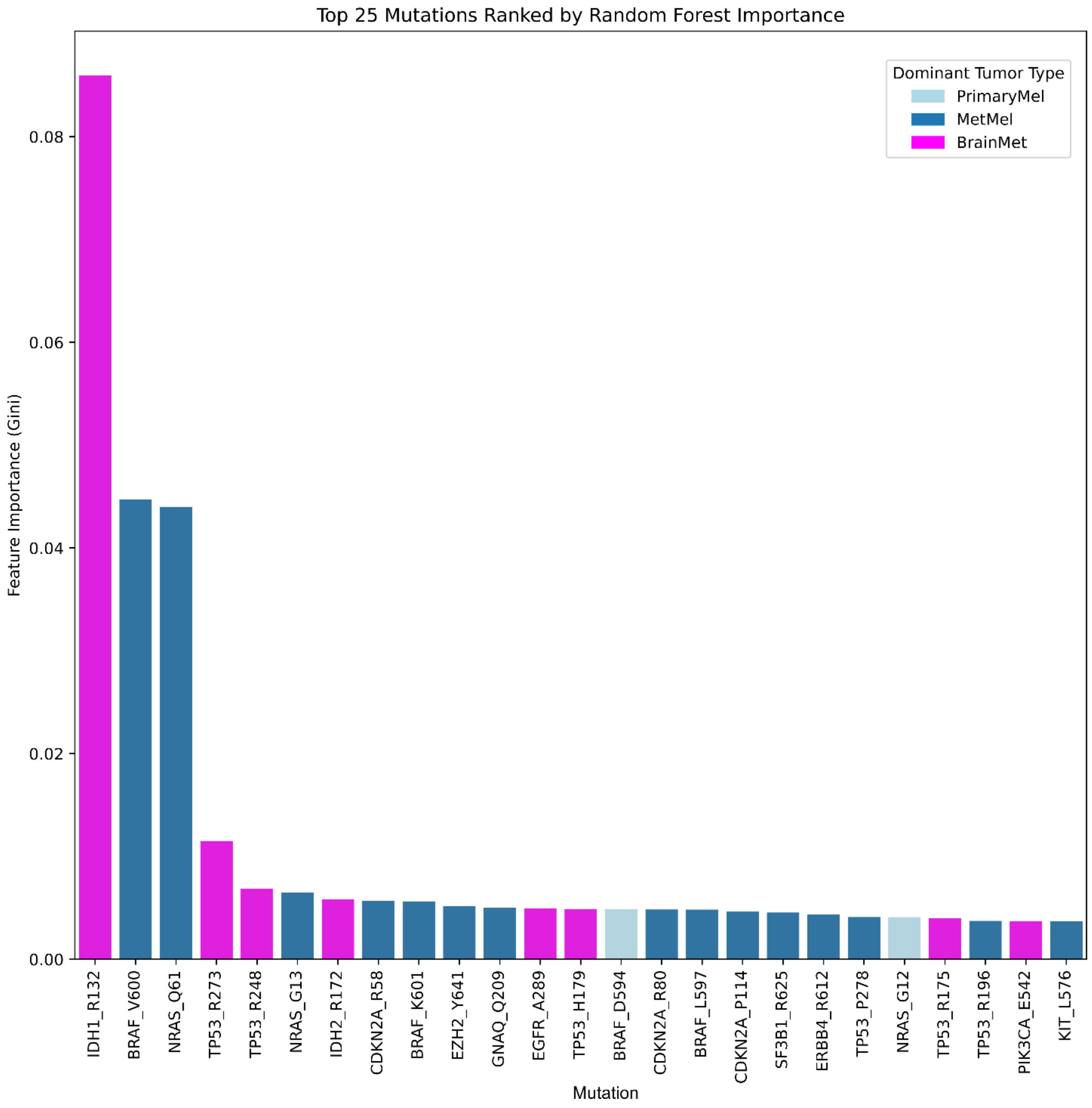


**Figure S3. Random forest classifier trained on binary gene mutation profiles to distinguish between primary melanoma, metastatic melanoma, and brain metastases.** Classification on the held-out test set showed highest precision and recall for metastatic melanoma (precision = 0.52, recall = 0.91, F1 = 0.66; n = 116), followed by brain metastases (precision = 0.89, recall = 0.43, F1 = 0.58; n = 77), and lowest performance for primary melanoma (precision = 0.24, recall = 0.06, F1 = 0.10; n = 62), with overall accuracy of 56% (n = 255). Feature importance, measured by the Gini index, identifies the top 25 genes contributing most to tumor type classification. Higher Gini values indicate greater contribution to reducing classification uncertainty across all trees, highlighting key mutational drivers distinguishing primary melanoma, metastatic melanoma, and brain metastases. Importance values are relative and reflect feature ranking rather than direction of effect.

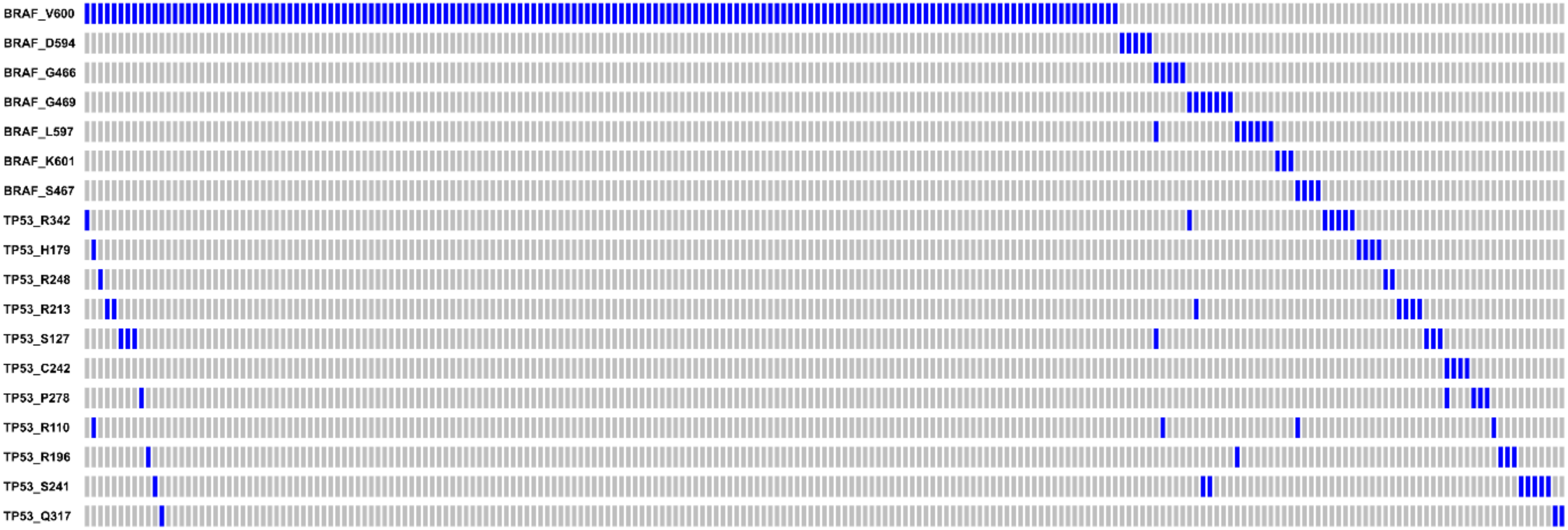
BRAF_V600
BRAF_D594
BRAF_G466
BRAF_G469
BRAF_L597
BRAF_K601
BRAF_S467
TP53_R342
TP53_H179
TP53_R248
TP53_R213
TP53_S127
TP53_C242
TP53_P278
TP53_R110
TP53_R196
TP53_S241
TP53_Q317

**Figure S4. Mutually exclusive BRAF and TP53 mutations in metastatic melanoma.** *BRAF*$^{V600}$ is the most frequent alteration, followed by non-V600 variants (D594, G466, G469, L597/K601, S467). BRAF mutations fall into three functional classes: Class I (V600) are strongly activating, RAS-independent monomers; Class II (G469, L597, K601) are RAS-independent dimers with intermediate-to-high activity; Class III (D594, G466, S467) are kinase-impaired but enhance RAS-dependent *CRAF* signaling. *TP53* mutations (R342, H179, R248, etc.) occur mutually exclusively with *BRAF*, likely reflecting redundant oncogenic roles: *BRAF* drives MAPK-mediated proliferation, whereas *TP53* loss promotes tumor progression via cell cycle, apoptosis, and genomic instability. This exclusivity indicates alternative oncogenic routes, with BRAF conferring proliferative signaling and *TP53* conferring resistance to stress and apoptosis.

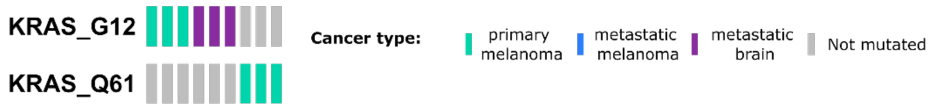


**Figure S5: Mutations in KRAS.** *KRAS*$^{G12}$ is observed in three primary melanoma and three metastatic brain samples, *KRAS*$^{Q61}$ is observed in three primary melanoma tumors.

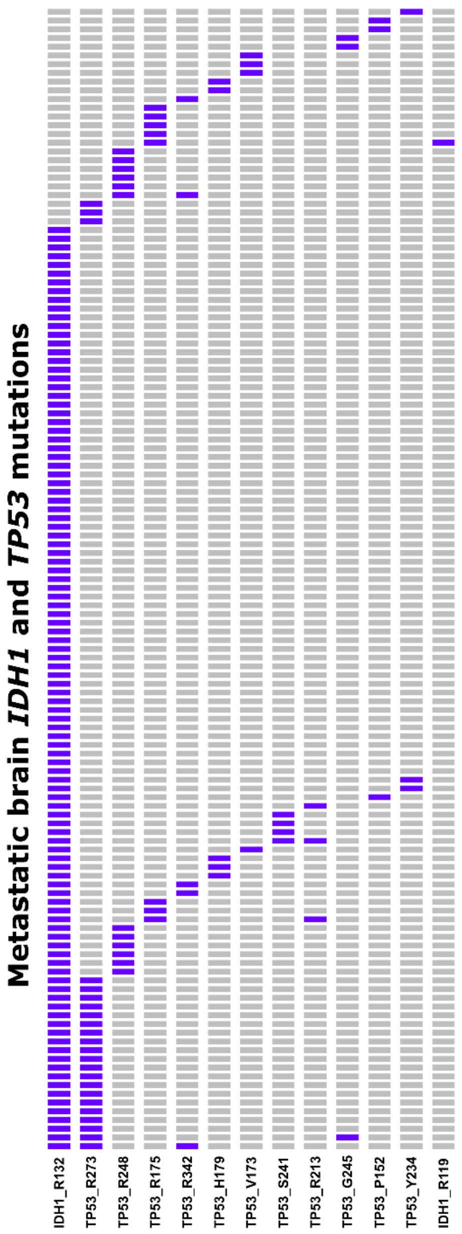


**Figure S6. Co-occurrence of IDH1 and TP53 mutations in metastatic brain tumors.** Columns represent individual tumors; rows denote recurrent mutations in *IDH1* or *TP53*. Purple bars indicate mutations. The canonical *IDH1*$^{R132}$ mutation is present in nearly all tumors, while $^{TP53}$ mutations span multiple hotspots (R273, R248, R175, R342, H179, S241, R213, etc.).

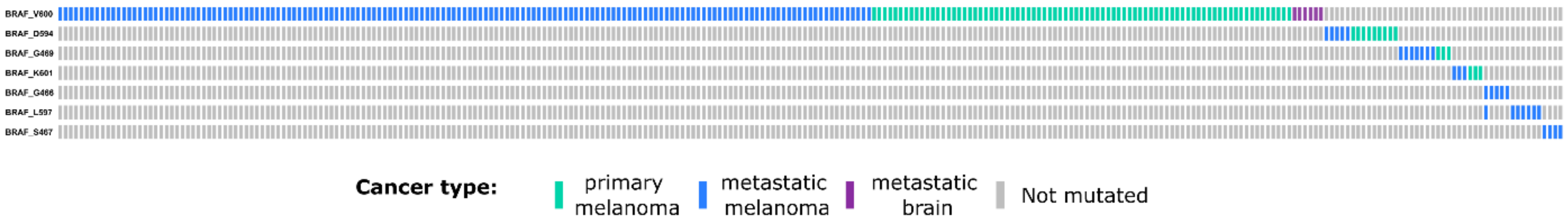
BRAF_V600
BRAF_D594
BRAF_G469
BRAF_K601
BRAF_G466
BRAF_L597
BRAF_S467
Cancer type:
primary melanoma
metastatic melanoma
metastatic brain
Not mutated

**Figure S7. Distribution of *BRAF* mutations across primary melanoma, metastatic melanoma, and metastatic brain tumors.** Each column represents an individual tumor, and each row corresponds to a recurrent *BRAF* mutation. Colored bars denote tumor type: green for primary melanoma, blue for metastatic melanoma, purple for metastatic brain tumors, and gray for samples without mutation. $BRAF^{V600}$ mutations were the most frequent and widely distributed across all tumor types, whereas non-V600 variants (D594, G466, G469, L597, S467) were less common and enriched in metastatic tumors.

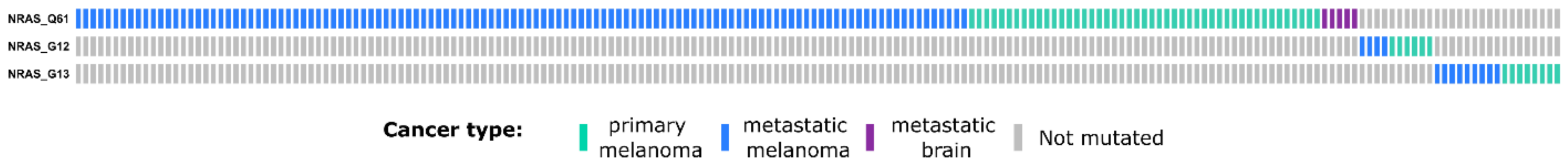

NRAS_Q61
NRAS_G12
NRAS_G13
Cancer type:
primary melanoma
metastatic melanoma
metastatic brain
Not mutated

**Figure S8. Distribution of *NRAS* mutations across primary melanoma, metastatic melanoma, and metastatic brain tumors.** Each column represents an individual tumor, and each row corresponds to a recurrent NRAS mutation. Colored bars denote tumor type: green for primary melanoma, blue for metastatic melanoma, purple for metastatic brain tumors, and gray for samples without mutation. $NRAS^{Q61}$ was the most frequent and widely distributed mutation across all tumor types, whereas $NRAS^{G12}$ and $NRAS^{G13}$ occurred less frequently and were more often observed in metastatic lesions.
